# Cerebrovascular Imaging-to-Graph Reconstruction for Individualized Digital Twin Brains

**DOI:** 10.64898/2026.06.24.734391

**Authors:** Chen Xie, Beini Hu, Abdulmalik M. Alakeel, Candace C. Fleischer, Andrei G. Fedorov

**Affiliations:** Woodruff School of Mechanical Engineering, Georgia Institute of Technology, Atlanta, GA, 30332, USA; Walter H Coulter Department of Biomedical Engineering, Georgia Institute of Technology and Emory University, Atlanta, GA, 30322, USA; Department of Radiology and Imaging Sciences, Emory University School of Medicine, Atlanta, GA, 30322, USA; Parker H. Petit Institute for Bioengineering and Bioscience, Georgia Institute of Technology, Atlanta, GA, 30332, USA

**Keywords:** Digital twin brain, Cerebrovascular graph reconstruction, Brain biophysical simulation

## Abstract

The development of digital twins in medicine, i.e., virtual replicas of human organs, offers a promising path toward precision medicine by enabling interpretable, mechanistic, and actionable insights. In the brain, cerebrovascular twins support individualized modeling of hemodynamics and bio-transport, with broad applications. A major bottleneck, however, is the lack of robust methods to transform *in vivo* cerebrovascular images into simulation-ready cerebrovascular meshes or graphs. Here, we present CerebroVascular Imaging to Graph reconstruction (CVIG), a robust and multiscale framework for reconstructing whole brain cerebrovascular graphs from *in vivo* cerebrovascular images. CVIG integrates vessel vectorization, with tolerance to discontinuity in vessel structures, using a topology-guided assembly of vessel trees to generate cerebrovascular graphs from medical images. We demonstrate the ability of CVIG to generate vascular graphs with improved vascular coverage and topological correctness, the capability essential for high fidelity brain biophysical simulations. This work establishes a vascular graph framework for individualized modeling and analysis, providing a key foundation for digital twins of the human brain.

## INTRODUCTION

Biomedical digital twins are patient-specific, data-assimilating, and dynamically evolving virtual replicas that aim to deliver mechanistic interpretation, prediction, and individualized decision support. [1–5] Digital twin brains (DTBs) are particularly compelling for precision medicine as they link individualized anatomy, biophysics, and physiology across scales within an interpretable and testable *in silico* framework, enabling computational prediction and simulation-based support for evaluating clinical scenarios and potential interventions.[6–8] Current work broadly follows two complementary directions. [1] First, DTBs can mimic local and whole-brain network dynamics by integrating structural and functional data to explain and forecast neural activity. [7,9,10] Second, physiological DTBs couple individualized anatomy with biophysical phenomena and regulation, most notably across the cerebral vasculature and cerebrospinal fluid, enabling quantitative modeling of hemodynamics and perfusion, [11–13] bio-transport, [14–17] solute clearance, [18–20] therapy delivery, [21–23] and hemodynamics based brain–machine interfaces (**Fig. 1**). [24] The increased interest in reliable DTBs stimulates new opportunities for both mechanistic brain research and clinical applications. [7,25,26]

**Figure 1.**
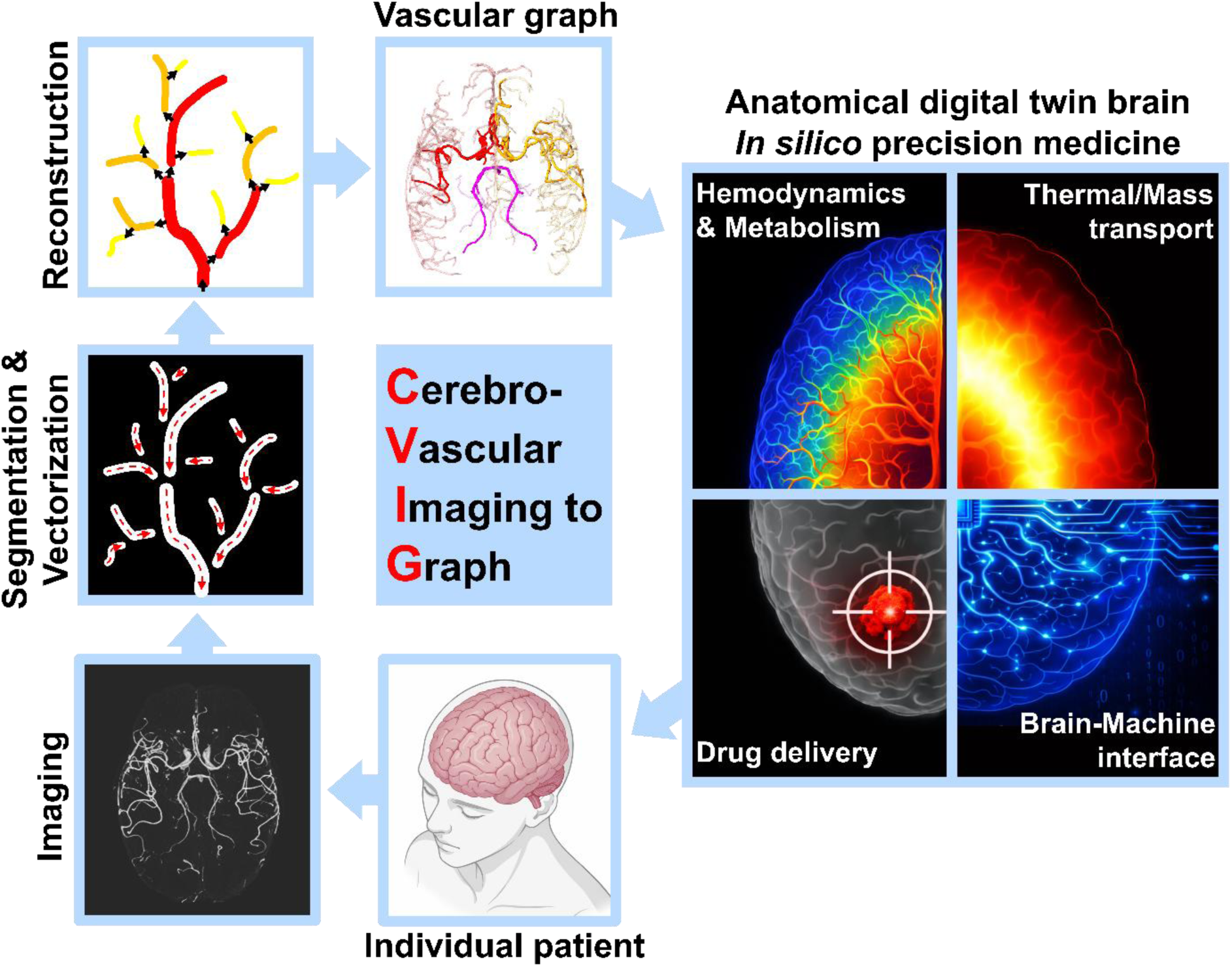
Schematic illustration of CerebroVascular Imaging to Graph (CVIG). CVIG comprises personalized cerebrovascular imaging, image processing and segmentation, vessel vectorization, and graph reconstruction. The resulting vascular graph provides an extensible foundation for anatomical digital twin brains (DTBs), with applications ranging from brain hemodynamics/perfusion, bio-transport and metabolism, and therapy delivery to brain–machine interfaces, enabling *in silico* precision medicine for individual patient.

Cerebrovascular twins, virtual anatomical replicas of the cerebral vasculature, are key building blocks of DTBs. [27–29] Current reconstruction pipelines generally involve two stages: pre-processing and segmentation of vascular images, [30,31] followed by construction of a simulation-ready mesh or graph; the latter remains to be a major bottleneck. [1,32,33] Considerable efforts have been devoted to reconstructing cerebrovascular twins. Linninger et al. have developed workflows to transform vascular imaging into structured meshes for hemodynamic modeling. [12,13,27,33,34] Chlebiej et al. proposed a customizable semi-automatic framework that combines vessel segmentation, skeleton optimization, and tubular reconstruction for 3D cerebrovascular modeling, reducing the need for extensive manual intervention. [35] Despite these advances, existing methods remain limited by their dependence on high quality imaging and segmentation. Recent studies have highlighted 1D vasculature graphs as promising substrates for multiscale/multi-physics brain modeling due to their flexibility and extensibility in large-scale multi-physical simulation. [15,28,36,37] Bartolo et al. developed a workflow from medical imaging to 1D graph construction for pulmonary and aortic vasculature. [38] Sung et al. extended this work to personalized cerebral vasculature maps and demonstrated the potential of 1D vascular graphs to facilitate whole brain hemodynamic and thermal modeling; [14,39–41] however, their pipeline relied on the Rivulet algorithm, originally designed for neuron tracking, and compatibility with complex multi-inlet cerebral vasculature of the human brain remains limited. [39,42,43] These gaps underscore the need for a robust methodology for personalized, simulation-ready cerebrovascular graph reconstruction from medical images acquired with multiple methods and of varied image quality.

We present CerebroVascular Imaging to Graph reconstruction (CVIG), an imaging-to-graph framework for robust reconstruction of personalized, whole brain cerebrovascular graphs from magnetic resonance angiography (MRA). As illustrated in **Fig. 1**, CVIG combines vessel enhancement and segmentation with Rivulet-inspired vessel vectorization that is tolerant to discontinuities in vasculature, followed by flow direction (inlet-to-outlet) and topology-guided tree assembly, in which inlet connectivity, upstream–downstream hierarchy, branch continuity, and cross-inlet separation are used to assemble fragmented vessel paths into physiologically accurate vascular graphs. This methodology improves vascular coverage, radius estimation, and topological correctness, achieving an approximately 100% increase in total graph length relative to the previous method by Sung et al. CVIG is compatible with multiple imaging modalities and parameters including time-of-flight (TOF) MRA, magnetic resonance venography (MRV), and computed tomography angiography (CTA) across different image resolutions. Importantly, we highlight CVIG-derived graphs substantially influence downstream brain biophysical simulation. The improved topological resolution of cerebrovascular graph enables improved spatial resolution and accuracy for virtual vessel-occlusion simulations, delivering anatomically plausible brain thermal responses that are infeasible with previous vessel graphs. By developing CVIG, we expect this work will establish a powerful cerebrovascular reconstruction framework, enable advanced cerebrovascular studies, and lay a key foundation for the development of DTBs.

## RESULTS

### Image preprocessing, cerebrovascular segmentation and vectorization

CerebroVascular Imaging to Graph reconstruction (CVIG) starts with processing *in vivo* cerebrovascular imaging and uses magnetic resonance angiography (MRA) volumes as the primary modality. As shown in **Fig. 2A**, the raw MRA volume was preprocessed before vessel vectorization, including resampling and intensity normalization (**Fig. 2Ai**), vessel enhancement and skull stripping (**Fig. 2Aii**), and subsequent vessel segmentation (**Fig. 2Aiii**). Vessel-like structures were enhanced using multiscale Hessian-based filtering, followed by skull stripping using a T_1_-weighted image-based brain mask. Instead of applying the T_1_-derived brain mask as a binary constraint, a soft spatial weighting mask using distance-transform-based inward feathering was applied to reduce boundary artifacts. The skull-stripped and vessel-enhanced MRA volume was then segmented by intensity-threshold-based foreground extraction to generate a cerebrovascular mask. Importantly, this cerebrovascular mask was not required to be topologically continuous but instead served as a permissive vessel domain for subsequent vectorization.

**Figure 2.**
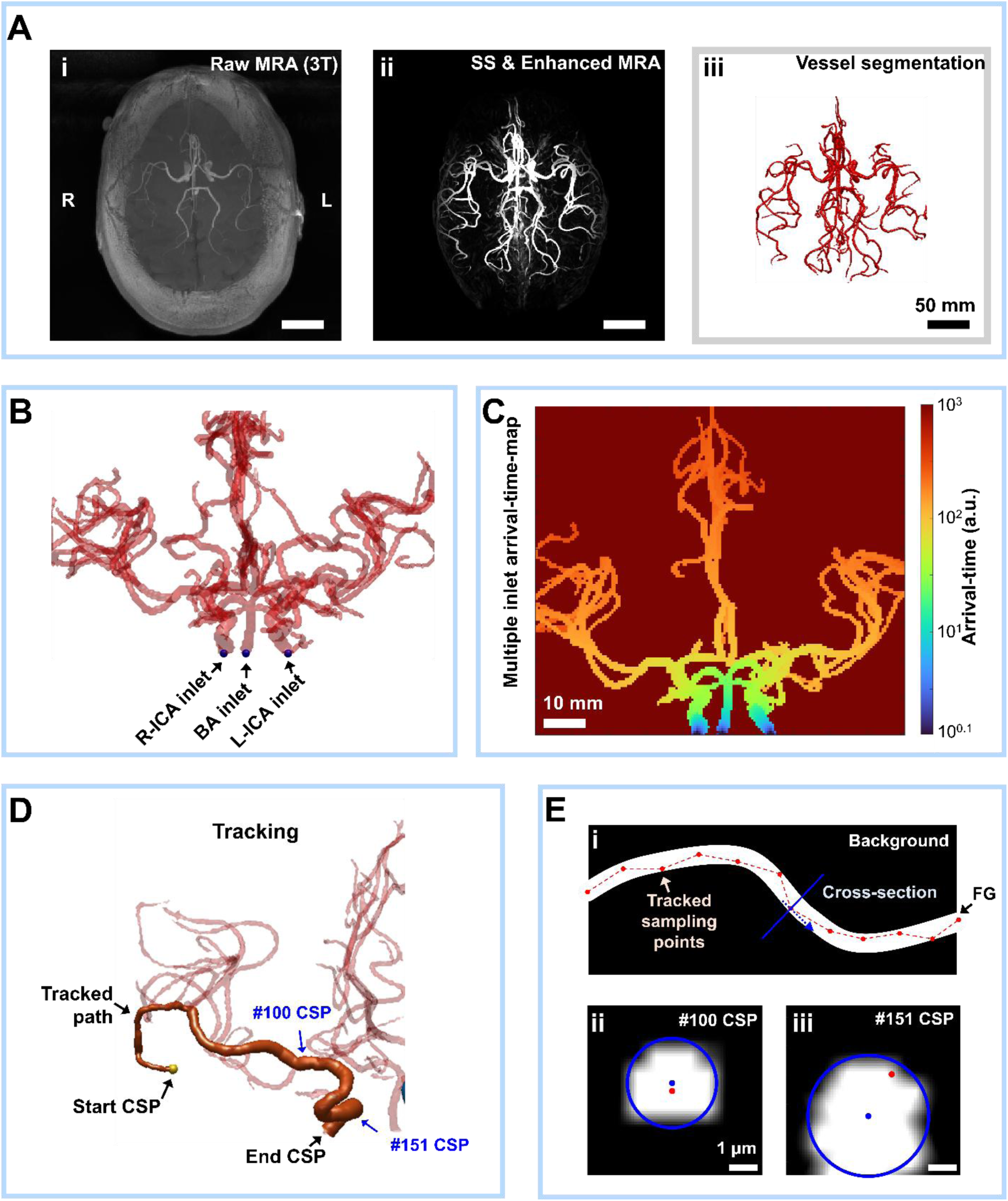
Methodology of Cerebrovascular (CV) segmentation and multi-inlet guided arrival-time-map solving and vessel tracking. (A) Key steps and outputs of MRA volume processing and segmentation, including resampling and normalization of the raw image (i), skull stripping (SS) using the T_1_-weighted image brain mask and denoising with multi-scale Frangi filter (ii), and segmentation using an intensity-based threshold (iii). (B) Multiple inlets of the cerebral artery tree at the bottom of the brain, the arrival-time-map corresponding to each inlet is estimated by solving the Eikonal equation. (C) Multi-inlet arrival-time-map is acquired by taking the voxel-wise minimum of arrival-time-maps for each inlet. (D) Vessel tracking starts from the centerline sampling point (CSP) with maximum arrival-time value within the foreground (FG). The vessel is tracked by discrete steepest descent on the arrival-time-map, following the locally smallest-valued neighboring voxel until reaching an inlet, a plateau, or a previously traced trunk. The highlighted #100 and #151 CSPs indicate representative locations along the tracked path used for the cross-sectional radius and centerline refinement examples shown in (E). (E) After the initial tracking (i), vessel radius is estimated and the centerline is refined by fitting inscribed circles along the traced path, with representative cross-sections shown at the #100 and #151 CSPs (ii &iii). Red and blue dots in ii & iii pinpoint initial and refined centerline, respectively; blue circles are cross sections of refined vectorized vessel.

Vessel vectorization was performed using a Rivulet-inspired tracking algorithm due to its strong tolerance to discontinuities and weak connectivity in tubular structures. [42,44] Notably, unlike the standard Rivulet algorithm, which typically starts from a single inlet, [42] CVIG adopts a multi-inlet strategy to reflect the physiological topology of the cerebral arterial system. As shown in **Fig. 2B**, three arterial inlets were automatically identified at the right internal carotid artery (R-ICA), left internal carotid artery (L-ICA), and basilar artery (BA). Compared with single-inlet tracking, the multi-inlet strategy provides a physiologically coherent guidance field for vessel tracking and further enables vessel-tree assembly using directionally guided blood flow (from inlets to outlet) and topological correction, such as to use of inlet identity and proximal-to-distal ordering to connect fragmented vessel paths while rejecting physiologically implausible cross-inlet or cross-branch connections.

For each inlet, an algorithmic arrival-time-map, representing the cumulative front-propagation cost from that inlet, was estimated over the segmented vessel domain by solving the Eikonal equation (Eqn. S1) numerically using a 3D multi-stencil fast marching method, [42,45–47] and a multi-inlet arrival-time-map was generated by taking the voxel-wise minimum arrival-time across all inlets (**Fig. 2C**, Eqn. S2). Here, a centerline sample point (CSP) denotes a discrete point along a traced vessel centerline that stores its spatial location and local vessel radius; each centerline is represented as an ordered sequence of connected CSPs from the tracking seed to the endpoint (**Fig. 2D&E**). Vessel tracking was initiated from the voxel with the highest arrival-time value in the segmented vessel domain, with an anatomy-guided weighting preference near the terminal regions of the anterior cerebral artery (ACA) and middle cerebral artery (MCA) to reduce unstable tracking from isolated noise voxels or weakly connected minor distal fragments, thereby preventing early tracking errors from propagating into the global topology of the reconstructed arterial tree (**Fig. 2D**). As shown in **Fig. 2C and 2D**, a downward tracking method is adopted and the tracking CSPs iteratively move to the neighboring voxel with the lowest local arrival-time value until reaching an inlet, or a previously traced trunk. The initially tracked voxel path was then refined by fitting the inscribed circle within the local vessel cross-section (**Fig. 2E**), where the fitted circle center was used to update the centerline location (**Fig. 2E ii&iii**, blue dot), while the vessel radius was estimated as the equivalent radius of the segmented cross-sectional area.

### Vessel-tree assembly using flow guidance and graph reconstruction

Next, the vectorized vessels, including tracked centerlines and local radius, were assembled into a topologically correct and physiologically accurate cerebrovascular tree. It should be noted that a flow direction–guided reconstruction strategy was adopted to encourage physiologically ordered expansion from proximal parent vessels toward distal child branches. In this strategy, inlet connectivity and distance from the inlets provide directional priors for inferring upstream–downstream hierarchy among CSPs, thus enabling physiologically consistent outward tree growth and reducing erroneous bridges between branches supplied by different inlets.

First, all tracked centerlines with radius were labeled as either inlet-connected or inlet-unconnected paths based on their connectivity to the arterial inlets (**Fig. 3A&B**). At the initial stage, only three root paths were inlet-connected, corresponding to the R-ICA, L-ICA, and BA. All remaining paths were treated as unconnected vessel segments to be evaluated during graph reconstruction.

**Figure 3.**
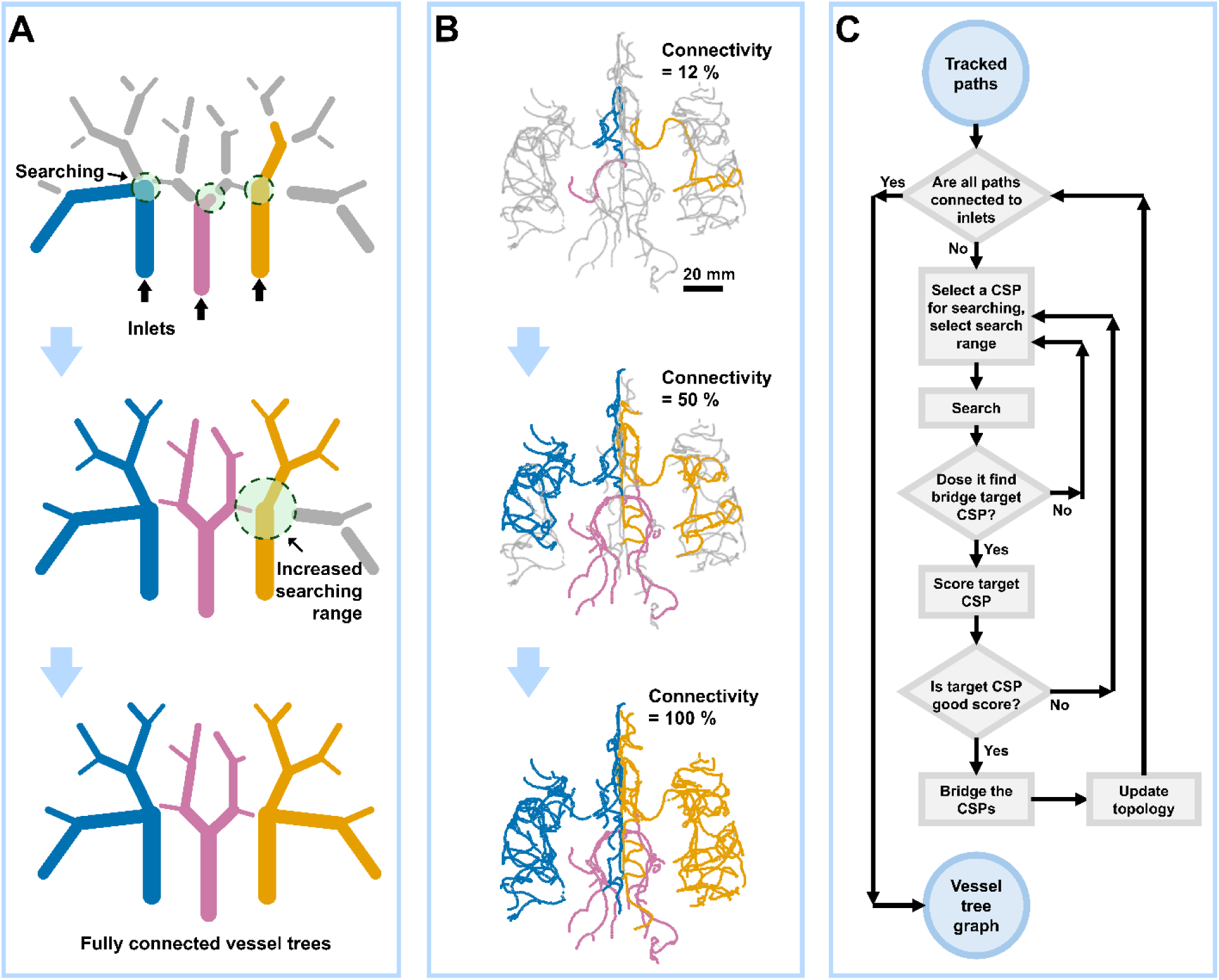
Methodology for image to graph reconstruction. (A) Schematic of inlet-guided cerebral artery tree reconstruction from tracked paths. Blue, purple, and orange paths are connected to the inlets, whereas gray paths are unconnected. Bridge searching starts from CSPs on connected paths; once a bridge is established, the vessel-tree topology is updated and all paths are relabeled. If the target connectivity level is not reached, the search is repeated with an expanded range (dotted circle) until the tree is fully connected. (B) Axial view of cerebral artery tree during reconstruction. Blue, purple, and orange paths are connected to the inlets (L-ICA, BA, R-ICA, respectively), whereas gray paths are unconnected. The reconstruction starts with minimal connectivity (12%) and ends with fully connected artery tree (connectivity = 100%). (C) Flowchart of the path-bridging and vessel tree graph reconstruction.

Second, to assemble the tracked centerlines into a complete and physiologically accurate vessel tree, CVIG expands the inlet-connected vessel tree by gradually connecting inlet-unconnected paths to inlet-connected paths. Here, a search–score–bridge strategy is adopted to connect the paths. Specifically, in each step, a CSP already belonging to the inlet-connected tree is selected and examined as the bridge-search origin CSP and used as the center of a spherical search region defined by a search range (**Fig. S1**). CSPs on nearby inlet-unconnected paths that fall within this spherical search region are identified as candidate bridge CSPs and evaluated as possible attachment points to the growing tree (**Fig. 3A&C and Fig. S1**). To ensure a physiologically ordered reconstruction, these search steps were guided by two priority rules (**Fig. S1**): a proximal-to-distal search order, in which CSPs closer to the inlet were selected as bridge-search origin CSPs before distal CSPs, and a CSP-type-dependent search range, in which special CSPs and terminal CSPs were assigned larger search ranges than ordinary centerline CSPs. Special CSPs were identified based on local geometric and topological features, including curvature and radius variation, as acute curvature or abrupt radius variation may indicate potential bifurcations or missed branch junctions.

Next, if candidate bridge CSPs were identified, then each of these candidate bridge CSPs was evaluated using a multi-score matching assessment composed of sub-scores ranging from 0 to 1 (**Figs. 3C and S2**, Eq. 1). These sub-scores included 1) diameter compatibility, which penalized or rejected connections from smaller upstream vessels to larger downstream vessels; 2) distance compatibility, which favored spatially closer target CSPs with least flow resistance connectivity; 3) directional consistency, which penalized abrupt angular changes between bridged paths as physically unrealistic; 4) connection smoothness, defined locally at the candidate bridge junction, which favored locally continuous transitions between the connected path, the bridge segment, and the unconnected path to form a vessel trajectory expected for hemodynamic flow, while penalizing abrupt kinks introduced by artificial bridging (**Fig. S2**). Additional anatomical constraint terms could also be included when needed, for example to prevent physiologically implausible assignments between left and right ACA branches. The total bridge score was defined as the product of all individual sub-scores:

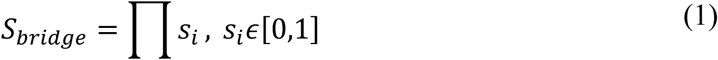

where *S_bridge_* is the total bridge score and *s_i_* represents each matching sub-score. Candidate bridges with any invalid matching term were rejected by assigning *S_bridge_* = 0. When multiple valid candidates were identified, the bridge with the highest total score was selected.

Once a candidate bridge CSP was accepted and the paths are bridged, the vessel-tree topology was updated and the newly connected path was relabeled according to its corresponding inlet branch. The newly formed attachment or bifurcation branches were then added back to the bridge-search origin CSP pool, allowing the connected tree to expand progressively toward distal branches (**Fig. 3B&C**). If the target connectivity, defined as the length fraction of inlet-connected paths over all tracked paths, was not reached after one search-score-bridge iteration, the global search range was increased and another search–score–bridge iteration was performed (**Fig. 3C**). It should be noted that the acceptance threshold for *S_bridge_* was progressively relaxed in later iterations to recover weakly connected or distal vessel fragments. After a predefined maximum number of strict scoring iterations, the algorithm entered a fallback connection mode in which the best available candidate within the search region was accepted to ensure full graph connectivity.This iterative reconstruction process converted the initially fragmented vectorized paths into a connected cerebrovascular graph that preserves centerline geometry, vessel radius, inlet assignment, and physiologically plausible branch topology for downstream biophysical simulations.

### CVIG improves cerebral vascular coverage, radius estimation and topological correctness

We systematically evaluated CVIG for cerebral artery graph reconstruction using images acquired from nine healthy human volunteers and one patient. The performance was assessed from two perspectives: vascular coverage, which measures the anatomical completeness of vascular structures recovered from the input image, and topological correctness, i.e., preservation of anatomical branch connectivity and upstream–downstream relationships in the reconstructed graph, which is essential as these relationships govern how flow, perfusion, heat, and other biophysical processes are supported by the cerebrovascular network. [48,49]

We first compared reconstructed cerebral artery trees across nine 3T TOF-MRA cases using three pipelines (**Fig. 4A–C**): a previous VesSeg + Rivulet method described by Sung et al.; [14,39,40] a mixed method using CVIG vessel segmentation followed by the original Rivulet tracking algorithm described by Liu et al., [39,42,43] without CVIG-specific multi-inlet guidance or topology-guided tree assembly; and the complete CVIG pipeline. As highlighted in **Fig. 4B**, CVIG achieved substantially improved vascular coverage, with an approximately 100% increase in total reconstructed vessel length across 3T cases (n = 9) relative to the Sung et al. method. This demonstrates its improved capability to recover vascular structures from *in vivo* MRA signals that were missed or underrepresented by prior reconstruction pipelines. Notably, the mixed CV + Rivulet method improved vascular coverage relative to the Sung et al. method but remained less complete than full CVIG, indicating that both improved vessel segmentation and the multi-inlet-guided graph reconstruction strategy contributed to the best outcome.

**Figure 4.**
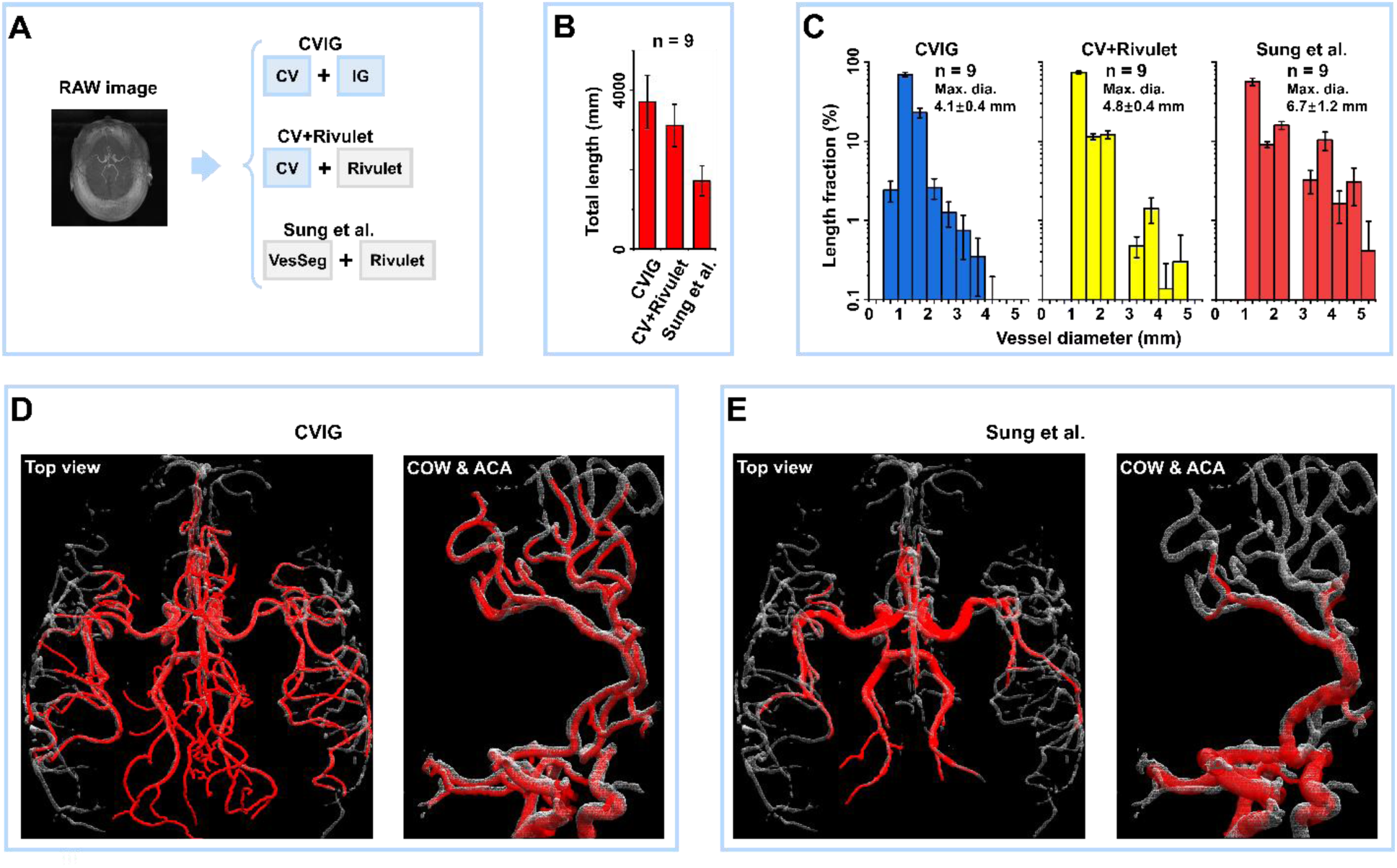
Validation of CVIG using 3T TOF-MRA and 7T anatomical reference. (A) Schematic of the reconstruction pipelines applied to 3T TOF-MRA data. The 3T image was processed using three methods: the complete CVIG, a mixed CV + Rivulet method using CVIG vessel segmentation followed by the original Rivulet tracking algorithm, and the previous VesSeg + Rivulet method (Sung et al). (B) Comparison of the total reconstructed cerebral artery tree length among the three methods across 3T TOF-MRA cases (n = 9). (C) Comparison of vessel diameter distributions of the reconstructed cerebral artery trees among the three methods across 3T TOF-MRA cases (n = 9). (D) Validation using paired 3T and 7T TOF-MRA from the same healthy volunteer. The cerebral artery graph reconstructed from 3T TOF-MRA using CVIG is shown as red tubes and overlaid on the 7T TOF-MRA-based vessel mask shown as a white mesh. (E) Reconstruction using the previous VesSeg + Rivulet method (Sung et al). The reconstructed graph is shown as red tubes and overlaid on the same 7T TOF-MRA-derived vessel mask shown as a white mesh.

Additionally, diameter distribution analysis showed CVIG produced more physiologically plausible vessel dimension predictions (**Fig. 4C**). The maximum vessel diameter estimated by CVIG was approximately 4–5 mm, comparable to that obtained by the mixed CV + Rivulet method and consistent with the expected size of proximal inlet arteries, [50] whereas the method described by Sung et al. yielded larger maximum diameters. Moreover, CVIG generated a smooth unimodal diameter distribution, while the reconstructions using the mixed method (CV + Rivulet) and the method described by Sung et al. showed more irregular bimodal distributions, indicating less stable radius estimation and discrepancy with physiologically expected results.

To further assess graph-to-anatomy agreement and topological correctness, we used a representative paired 3T/7T TOF-MRA dataset from the same healthy volunteer, with the higher-resolution 7T MRA (voxel size = 0.3 × 0.3 × 0.5 mm) serving as an anatomical benchmark reference (**Fig. 4D&E**). As illustrated in **Fig. 4D&E**, the reconstructed vascular graphs, shown as red tubes, were overlaid on the 7T TOF-MRA-derived vessel mask, shown as a white mesh, to visually evaluate their agreement with the underlying vascular anatomy. In the Circle of Willis (CoW) and anterior cerebral artery (ACA) regions, CVIG improved vascular coverage while correctly resolving the bilateral ACA branches. In contrast, the previous VesSeg + Rivulet method (Sung et al.) produced a coarse, enlarged structure that merged the two ACA branches. These results highlight the importance of evaluating not only vascular coverage but also the topological correctness of reconstructed cerebrovascular graphs.

Next, we evaluated the performance of CVIG across MRA volumes with different spatial resolutions. As shown in **Fig. S3A**, a high-resolution 7T MRA volume was resampled to generate high-resolution (voxel size = 0.3 × 0.3 × 0.3 mm^3^), medium-resolution (0.6 × 0.6 × 0.6 mm^3^), and low-resolution (1.2 × 1.2 × 1.2 mm^3^) datasets. The high-resolution reconstruction, followed by extensive manual correction, was used as the reference vascular graph, while the medium- and low-resolution volumes were processed using CVIG, the Sung et al. method, and the mixed CV + Rivulet method. This controlled design of target datasets preserved the same underlying vascular anatomy while systematically degrading image resolution for rigorous comparative assessment.

Across medium- and low-resolutions, CVIG achieved the highest centerline coverage among all three methods at every tested resolution (**Fig. S3B**, Eqn. S3). Although the mixed CV + Rivulet method achieved comparable but slightly lower centerline coverage, CVIG showed a significant advantage in topological correctness of the vascular graph. We therefore focused on the ACA as a representative anatomically complex branch that is important for resolving cerebral arterial territories, yet remains challenging to reconstruct accurately from *in vivo* vascular images. [28,51] Demonstrated in **Fig. S3C**, the reconstructed R-ACA (red) and L-ACA (blue) branches in the reference graph were clearly separated. Qualitative comparison showed that CVIG significantly improved the preservation of the bilateral ACA topology across all resolutions, whereas the other reconstructions showed major topological errors, including missing branches, incorrect cross-branch connections, and merging of the two ACA branches (**Fig. S3D**). Quantitative analysis of true, false-negative, and false-positive ACA length fractions further confirmed that CVIG consistently preserved a larger fraction of correct ACA topology and reduced erroneous reconstruction compared with the mixed CV + Rivulet and Sung et al. methods (**Fig. S3E&F**). Together, these results demonstrate that CVIG improves both vascular coverage and anatomical topology preservation across varying MRA spatial resolutions.

### CVIG performs robustly across diverse cerebrovascular imaging inputs

Given the improved performance of CVIG as applied to TOF-MRA input images, we further demonstrated that CVIG can robustly reconstruct physiologically coherent cerebrovascular graphs for other vascular imaging inputs, including magnetic resonance venography (MRV) and computed tomography angiography (CTA). As shown in **Fig. 5A–C**, CVIG reconstructed a cerebral venous graph from 3T MRV data despite its lower through-plane resolution and different vascular contrast relative to TOF-MRA, enabling reconstruction of both cerebral arteries and veins in the same subject (**Fig. 5B**). In addition, CVIG was applied to CTA, where it reconstructed a cerebral artery graph from CTA-derived vascular signals (**Fig. 5D–F**). These results demonstrate that CVIG is not limited to MRA processing but can serve as a general cerebrovascular imaging-to-graph framework across arterial and venous imaging modalities.

**Figure 5.**
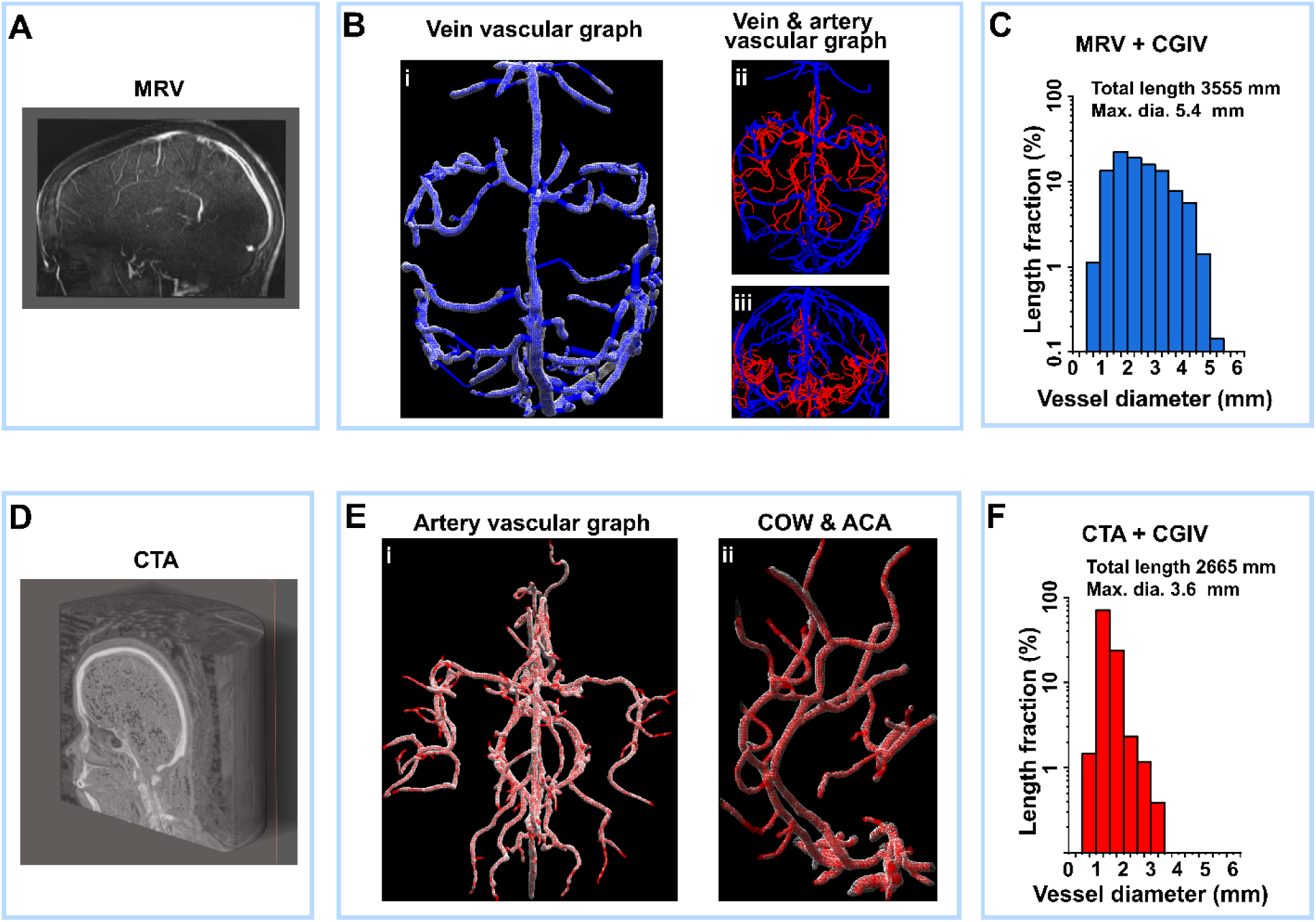
Generalizability of CVIG to MRV and CTA for cerebrovascular graph reconstruction. (A) Raw MRV image (voxel size = 0.8 × 0.8 × 2 mm^3^). (B) CVIG reconstruction of the cerebral vein graph from MRV, blue tube indicating reconstructed vein tree; white mesh (i) demonstrate MRV vessel mask, and red tubes are reconstructed artery tree for same patient (ii & iii). (C) Diameter distribution of the reconstructed vein tree from MRV. (D) Raw CTA image (voxel size = 0.5 × 0.5 × 0.6 mm^3^). (E) CVIG reconstruction of the cerebral artery graph from CTA, red tube indicating reconstructed artery tree, white mesh demonstrates vessel mask. (F) Diameter distribution of the reconstructed artery tree from CTA.

### CVIG-derived vascular graphs enable improved spatial resolution and accuracy of vessel occlusion simulations in brain thermal response

One of the key objectives for digital twins is to deliver predictive, interpretive, and first principle-based modeling results with virtual experiments that are otherwise impractical, costly, or impossible to perform. Here we demonstrate that the improved topological resolution of CVIG-derived cerebrovascular graphs significantly affects downstream brain biophysical simulation, enabling virtual cerebrovascular occlusion simulations with much improved resolution and anatomically plausible thermal responses that were not achievable with previous vascular reconstructions. As shown in **Fig. S4**, the vessel tree graphs reconstructed by the Sung et al. method and by CVIG were incorporated into a brain thermal transport model to simulate whole brain temperature. [41] It should be noted that, even though the default simulated temperature maps using two vessel tree graphs appear similar due to the temperature smoothing effect of heat conduction, results from virtual vessel-occlusion simulations demonstrate remarkable contrast. As illustrated in **Fig. 6A and 6B**, using the vessel tree generated by the Sung et al. method, the ACA region could not be selectively separated into anatomically distinct unilateral branches (**Fig. 4D, S3C and S3D**). As a result, the left and right ACA branches could not be independently targeted in the Sung et al. vessel tree. The virtual occlusion therefore had to be applied to the merged ACA-associated structure, leading to a large and non-specific bilateral thermal perturbation (artificially higher temperature due to lack of cooling by arterial blood) in the simulated brain temperature map (**Fig. 6B**). On the other hand, the higher topological resolution of the CVIG-derived graph enabled simulation of selective virtual occlusion of individual ACA branches (**Fig. 4E, S3C and S3D**). When the R-ACA branch was selectively blocked (**Fig. 6C**), the resulting temperature perturbation was spatially localized to the corresponding vascular territory rather than spreading broadly across both hemispheres (**Fig. 6D**). A complementary left ACA blockage experiment produced a side-specific thermal response on the opposite side, further supporting the anatomical specificity and its importance to predicting an accurate temperature response (**Fig. S5**). Together, these results demonstrate that CVIG-derived cerebrovascular graphs do more than improve vessel visualization or reconstruction metrics: they enable branch-specific, anatomy-aware what-if simulations using a physics-based brain biophysical model, providing a concrete step toward realizing individualized DTBs.

**Figure 6.**
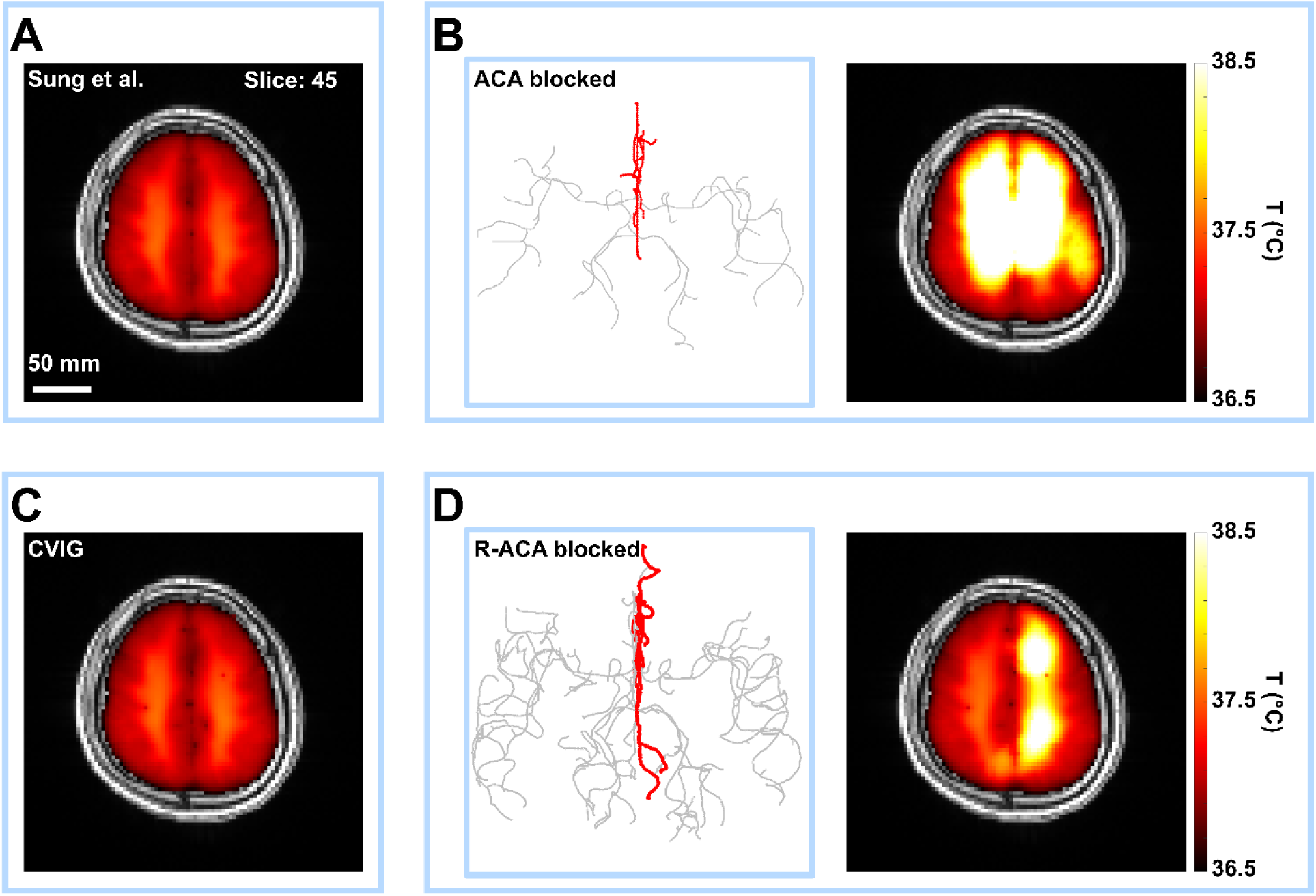
CVIG enables improved resolution and accuracy for virtual vessel-occlusion simulations in brain thermal transport modeling. (A) Baseline simulated brain temperature map with the vessel tree reconstructed by the Sung et al. method. Representative slice is shown. (B) Virtual occlusion scenario using the Sung et al. vessel tree, in which the ACA-associated branches were blocked. Red curves indicate the blocked vessel branches. (C) Baseline simulated brain temperature map using the vessel tree reconstructed by CVIG. (D) Virtual occlusion scenario using the CVIG-derived vessel tree, in which the right anterior cerebral artery (R-ACA) branch was selectively blocked. Red curves indicate the blocked vessel branches. The higher topological resolution of the CVIG-derived graph enables branch-specific perturbation and produces a localized thermal response in the corresponding vascular territory.

## DISCUSSION

We developed a high fidelity imaging-to-graph framework CVIG for reconstructing simulation-ready cerebrovascular graphs. We demonstrated it improved vascular coverage and enhanced graph topological correctness. This improved performance arises from two key design features. First, CVIG uses vectorization that is tolerant to discontinuity within a permissive vessel domain, allowing weakly connected or fragmented vascular signals to be retained rather than discarded during early processing, thereby improving vascular coverage. Second, CVIG uses topology-guided vessel-tree assembly strategy instead of purely local centerline connection approach. By incorporating topological factors, including inlet identity, proximal-to-distal ordering, diameter compatibility, and vessel-path smoothness, CVIG connects fragmented vessel paths based on physiological rules, reducing erroneous cross-branch or cross-territory connections and improving graph topological correctness. Although CVIG was specifically developed for cerebrovascular reconstruction, its strategy may provide a generalizable framework for constructing vasculature graphs in other organ-specific digital twins.

Despite these substantial advances, additional enhancements can be recommended. First, CVIG is sensitive to correct inlet identification and initial (near-inlet branches) reconstruction of major inlet-connected vessels. Since downstream graph assembly is guided by flow direction guided inlet connectivity and upstream–downstream hierarchy, errors in inlet localization or proximal vessel tracking may propagate through the reconstruction and cause global topological errors. Future work should aim to improve inlet detection, introduce uncertainty-aware tracking, and incorporate anatomical checks during vessel vectorization. Second, multi-inlet arrival-time-map tracking requires solving the Eikonal equation, which can be computationally demanding for high-resolution volumetric images. Third, although the current search–score–bridge strategy improves topological correctness, graph assembly remains locally optimized and lacks a global topology check after assembly. Extension of CVIG approach could incorporate iterative whole-tree scoring after bridging to further refine reconstructed graphs.

Our results also highlight the continuing importance of high-quality vascular imaging. Although CVIG remained robust across MRA resolutions, resolution-degradation analysis showed that algorithmic reconstruction cannot fully recover vascular information absent from the input image (**Fig. S3**). This limitation affects both distal branches and major trunks when partial-volume effects, low signal-to-noise ratio, or discontinuous signals compromise vessel visibility. To address sub-resolution vasculature, future work could combine CVIG-derived individualized large-vessel graphs with statistically generated microvascular networks that preserve both population-level vascular architecture and subject-specific anatomy. [12,15,28,51,52]

## CONCLUSION

This study establishes CVIG as a powerful imaging-to-graph framework for reconstructing individualized cerebrovascular graphs from vascular images. The key result is that CVIG improves vascular coverage, radius estimation, and topological correctness over prior pipelines, while remaining robust across MRA resolutions and vascular imaging modalities. Importantly, we demonstrate that CVIG-derived graphs can significantly enhance downstream brain biophysical simulations, producing physiologically plausible virtual-experiment results not achievable with the current state-of-the-art methods. This work advances cerebrovascular reconstruction and provides a foundational capability for realizing individualized digital twin brains.

## METHOD AND MATERIAL

### Image acquisition

This study was approved by the Emory University Review Board. Retrospective MR data from nine healthy volunteers (3 females, 6 males; mean ± standard deviation (SD) age = 27 ± 3.5 years old) were used to develop the method. MR data in healthy volunteers were acquired with a Siemens 3T Prisma FIT whole body MR scanner (Siemens Healthineers, Erlangen, Germany) using a 32-channel head and neck coil (Siemens). In one subject, data were also acquired with a Siemens 7T Terra whole body MR scanner (Siemens) using a 8-channel transmit, 32-channel receive head and neck coil (Nova Medical, Wilmington, MA, USA).

For 3T acquisition, structural images were obtained using a T_1_-weighted magnetization prepared rapid gradient echo (MPRAGE) sequence (Repetition time(TR)/Inversion time(TI)/Echo time (TE)/Flip angle (FA)= 2300/900/3.39ms/9, matrix = 192 × 192, FOV = 256 × 256 mm², voxel size = 1.33 × 1.33 × 1.0 mm³, scan time = 4 min 8 s). At 7T, a T_1_-weighted MP2RAGE was used (TR/TI1/TI2/TE = 4300/1000/3200/2.19 ms, 1 mm isotropic resolution, scan time = 5 min 46 s).

Vascular imaging included 3D time-of-flight (TOF) magnetic resonance angiography (MRA) at 3T (TR/TE = 22/3.86 ms, flip angle = 15, slice thickness = 0.62 mm) with different sets of FOVs and matrix sizes (set 1: FOV = 200 × 200 mm², matrix = 256 × 256; set 2: FOV = 220 × 220 mm², matrix = 256 × 256;). At 7T, a 3D TOF MRA was acquired with the following parameters: TR/TE = 19/3.42 ms, flip angle = 24, slice thickness = 0.5 mm, set 1: FOV = 181 × 200 mm², matrix = 696 × 768.

To evaluate the capability of CVIG using multiple imaging modalities, MRV was also used from a subset of healthy volunteers (n=1).At 3T, 2D TOF MRV was acquired (set 1: TR/TE = 18/3.79 ms, flip angle = 60, FOV = 220 × 220 mm², matrix = 256 × 256, slice thickness = 3.0 mm; set 2: TR/TE = 19/3.79 ms, flip angle = 60, FOV = 220 × 198 mm², matrix = 256 × 230, slice thickness = 3.0 mm).

To evaluate multiple imaging modalities, clinical CT and MRI images acquired from a unilateral acute ischemic stroke patient (Female, Age = 49 years old) were also included. CT imaging was performed on a Discovery CT750 HD scanner (GE Healthcare). Imaging protocol included CTA (500 mA,120 kVp, resolution= 0.5 × 0.5 × 0.6 mm³) and CTP (145 mA, 80 kVp, 0.47 × 0.47 mm² pixel spacing). MRI data was acquired on a 3T TIM Trio scanner (Siemens Healthcare), including post-contrast T1-weighted MPRAGE (TR/TI/TE = 1900/900/2.52 ms, flip angle = 9, 1.0 mm isotropic).

### Image preprocessing and vessel segmentation

All simulations were performed using MATLAB (Version R2023a, Mathworks, Natick, MA). The MRA, T1 and MRV volumes first transformed into 3D matrixes and subsequently normalized and resampled (cubic interpolation). For 3T images, the resampled voxel size is 0.6 × 0.6 × 0.6 mm^3^, for 7T images, the resampled voxel sizes are: high-resolution (voxel size = 0.3 × 0.3 × 0.3 mm^3^); medium-resolution (0.6 × 0.6 × 0.6 mm^3^); and low-resolution (1.2 × 1.2 × 1.2 mm^3^);

The T1 and MRV volumes were co-registered to the MRA space using rigid transformation. The transformation was initialized by center-of-mass alignment, and followed by small-angle rotation search, and final axis-wise line search. Vessel-like structures in the MRA and MRV volumes were enhanced using the MATLAB Fibermetric function, which applies a Hessian-based multiscale Frangi vesselness filter to 3D grayscale volumes, with the pre-set diameter of 1-12 voxels. Skull stripping is performed with brain mask, generated based on T_1_-weighted image with ANFI 3dSkullStrip, Version AFNI_25.0.00. [53,54]

The skull-stripped and vessel-enhanced MRA volume was segmented using intensity-threshold-based foreground extraction, where voxels above the percentile threshold (default: 99%) were retained as the initial vessel foreground. The foreground mask was then refined by connected-component filtering (18-connectivity), in which connected components smaller than certain voxels were removed, followed by filling of internal cavities fully enclosed by the vessel foreground.

## ACKNOWLEDGEMENTS

Research reported in this publication was supported by the National Institutes of Health and the National Institute for Neurological Disorders and Stroke under award number R01NS138044.

## Supporting Information

**Figure S1.**
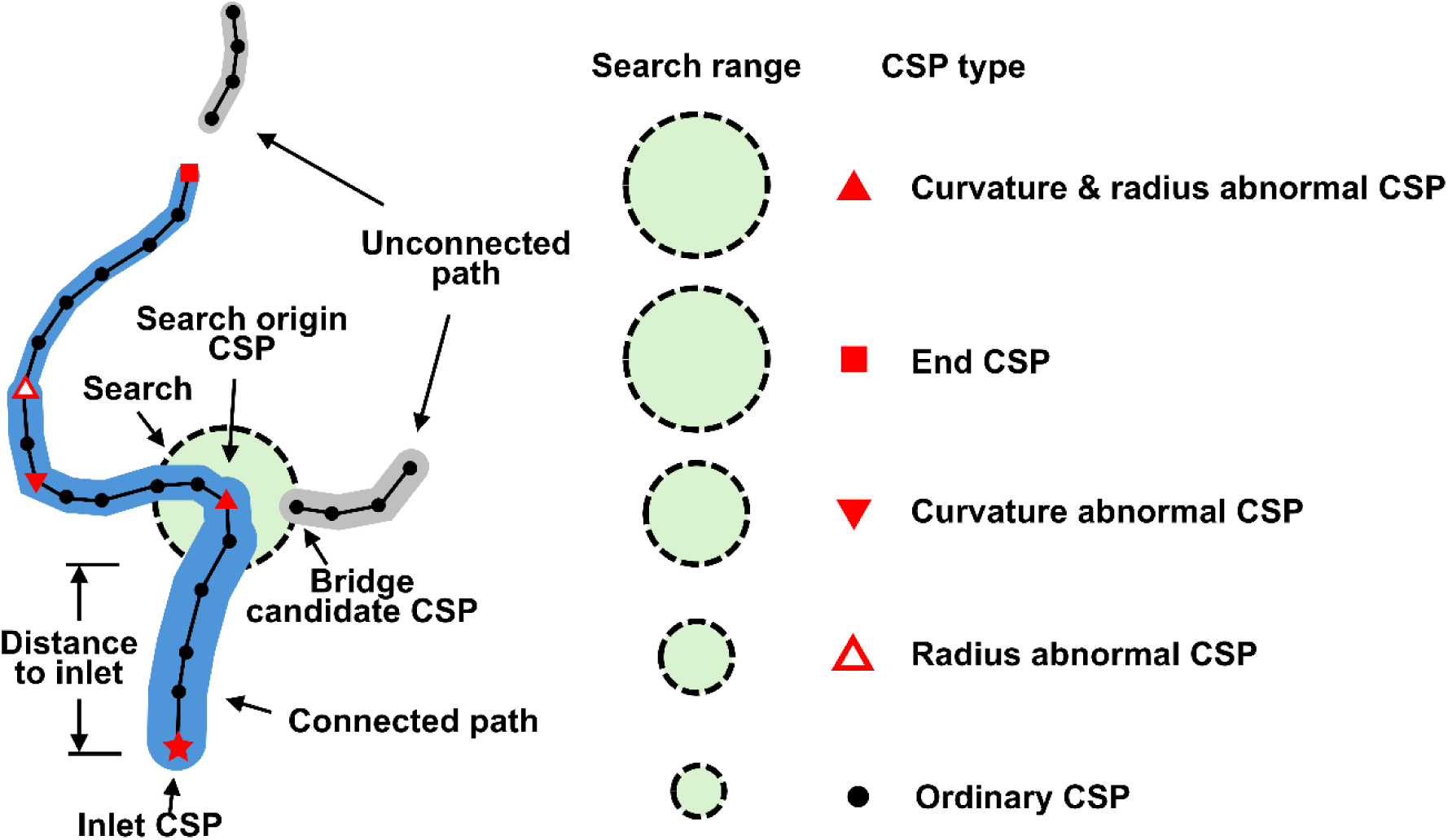
Schematic illustration of the search–bridge algorithm. Bridge searching starts from centerline sampling points (CSPs) on inlet-connected paths and proceeds according to two priority rules to ensure a physiologically ordered reconstruction. First, CSPs closer to the inlet are processed. Second, special CSPs are assigned with larger search ranges than ordinary CSPs. These special CSPs are identified and prioritized as potential search origin CSPs based on local geometric and topological features, including curvature, radius variation, and CSP type, such as terminal CSPs. CSPs from unconnected path that fall in the search range are then identified as bridge candidate CSPs.

**Figure S2.**
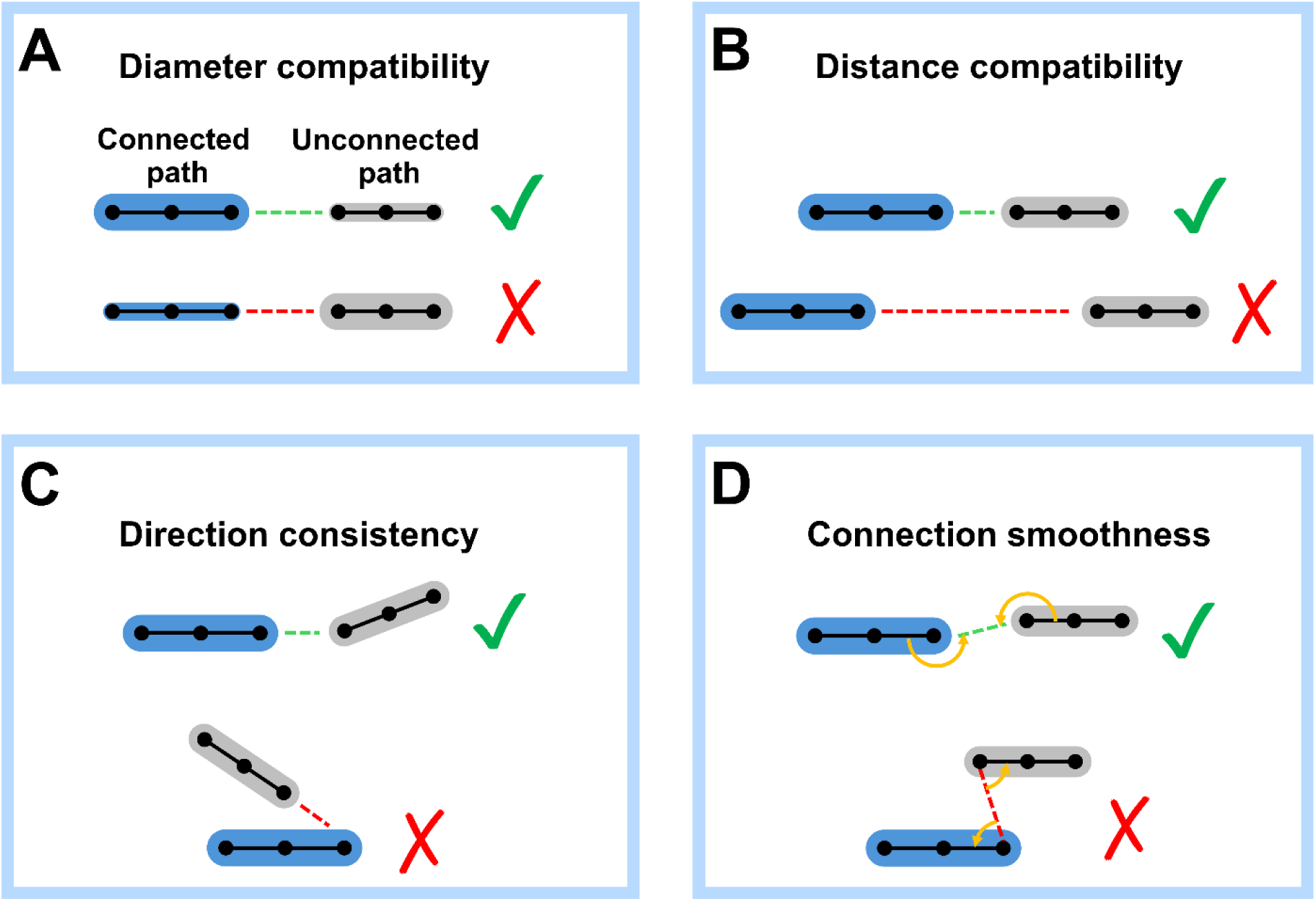
Schematic illustration of bridge-candidate matching scores. Candidate target CSPs are evaluated using geometry- and topology-aware scores before bridging. These include (A) diameter compatibility, where connections from smaller upstream vessels to larger downstream vessels are penalized or rejected; (B) distance compatibility, where closer target CSPs are preferred; (C) directional consistency, where abrupt angular changes between bridged paths are penalized or rejected; and (D) connection smoothness, where the local continuity introduced by the bridge is evaluated using the two junction angles formed between the connected path, the bridge segment, and the unconnected path. A fully smooth connection corresponds to junction angles approaching 180°, whereas lower scores are assigned when either angle approaches 90°; candidate bridges producing acute junction angles (< 60°) are rejected.

**Figure S3.**
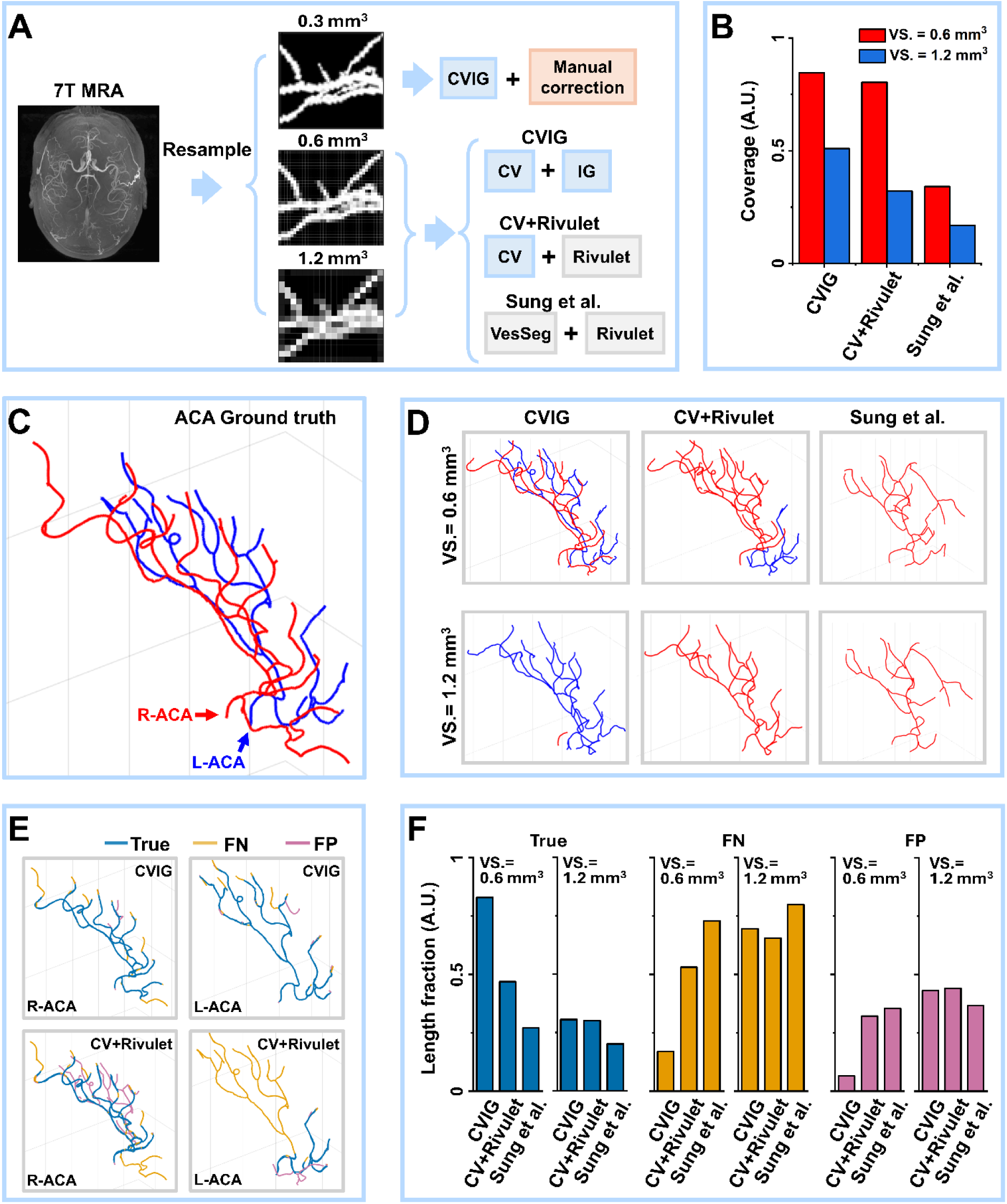
Vessel coverage and topological correctness of CVIG across various resolution of MRA images. (A) 7T MRA image (voxel size = 0.3 × 0.3 × 0.5 mm^3^) is resampled into high resolution image (voxel size = 0.3 × 0.3 × 0.3 mm^3^), medium resolution image (voxel size = 0.6 × 0.6 × 0.6 mm^3^), and low resolution image (voxel size = 1.2 × 1.2 × 1.2 mm^3^). The high resolution image then goes through CVIG and extensive manual correction, and the final result is taken as the ground truth. The medium and low resolution images are processed by CVIG, prior method using VesSeg + Rivulet (Sung et al.), and mixed method (CV + Rivulet). (B) Comparison of reconstructed vessel tree centerline coverage using all three methods. (C) The centerline of anterior cerebral artery (ACA) using ground truth vessel tree. (D) The centerline of ACA using all three methods on both medium- and low-resolution images. (E) Topological correctness of ACA from reconstructed cerebral artery trees by CVIG and mixed method for medium-resolution image. (F) Comparison of length fraction of correct (True), false negative (FN), false positive (FP) segments of ACA tree by all three methods for medium- and low-resolution images.

**Figure S4.**
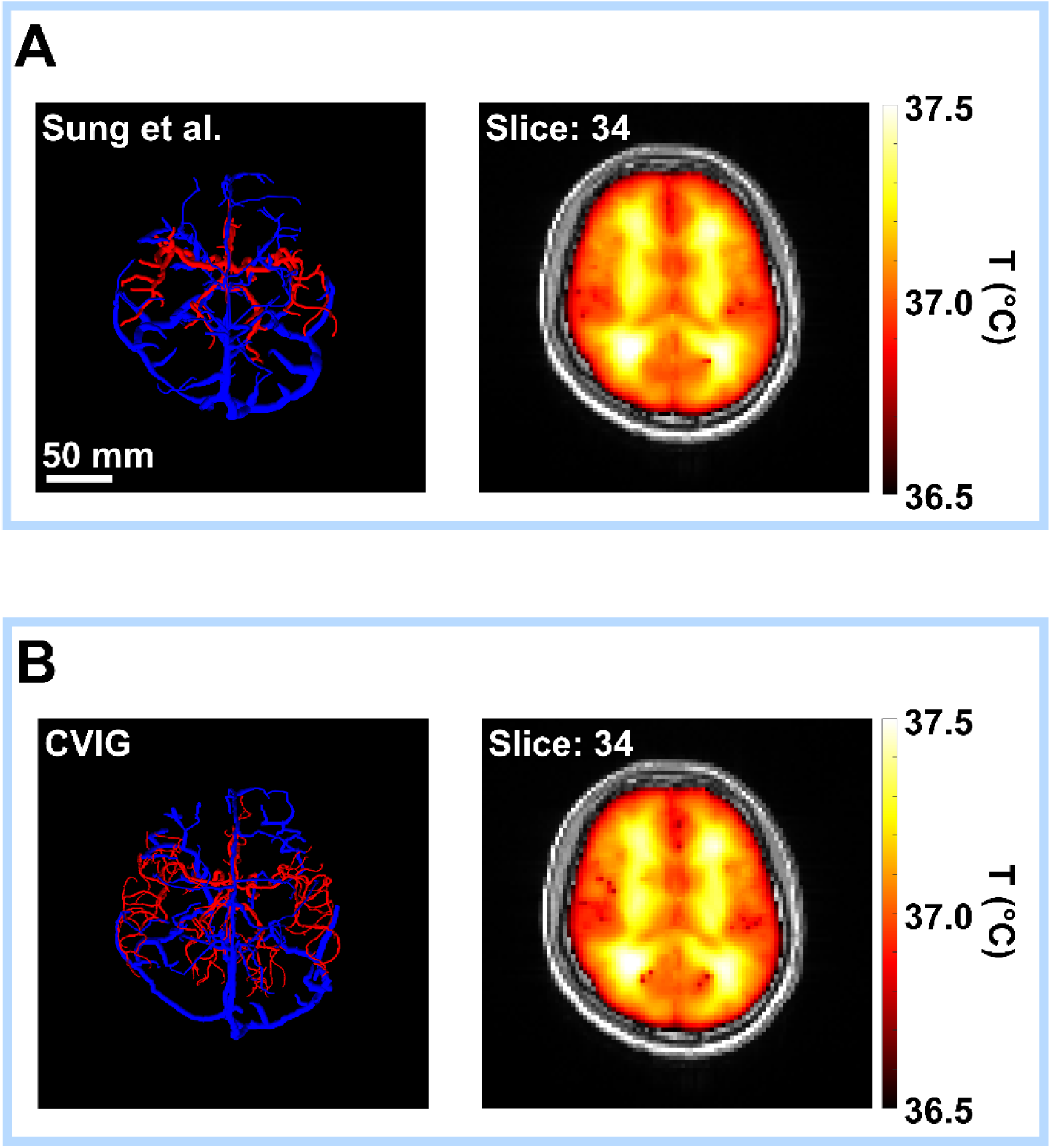
Baseline brain temperature predictions using vessel trees reconstructed by the Sung et al. method and CVIG. (A) Reconstructed artery and vein vessel trees using the Sung et al. method and the corresponding simulated baseline brain temperature map. Red and blue tubes indicate arterial and venous vessel trees, respectively. Total slide number: 80. (B) Reconstructed artery and vein vessel trees using CVIG and the corresponding simulated baseline brain temperature map. Compared with the Sung et al. reconstruction, CVIG provides a more spatially extensive and topologically resolved cerebrovascular graph while producing a comparable baseline temperature distribution under the non-occlusion (healthy vasculature) condition.

**Figure S5.**
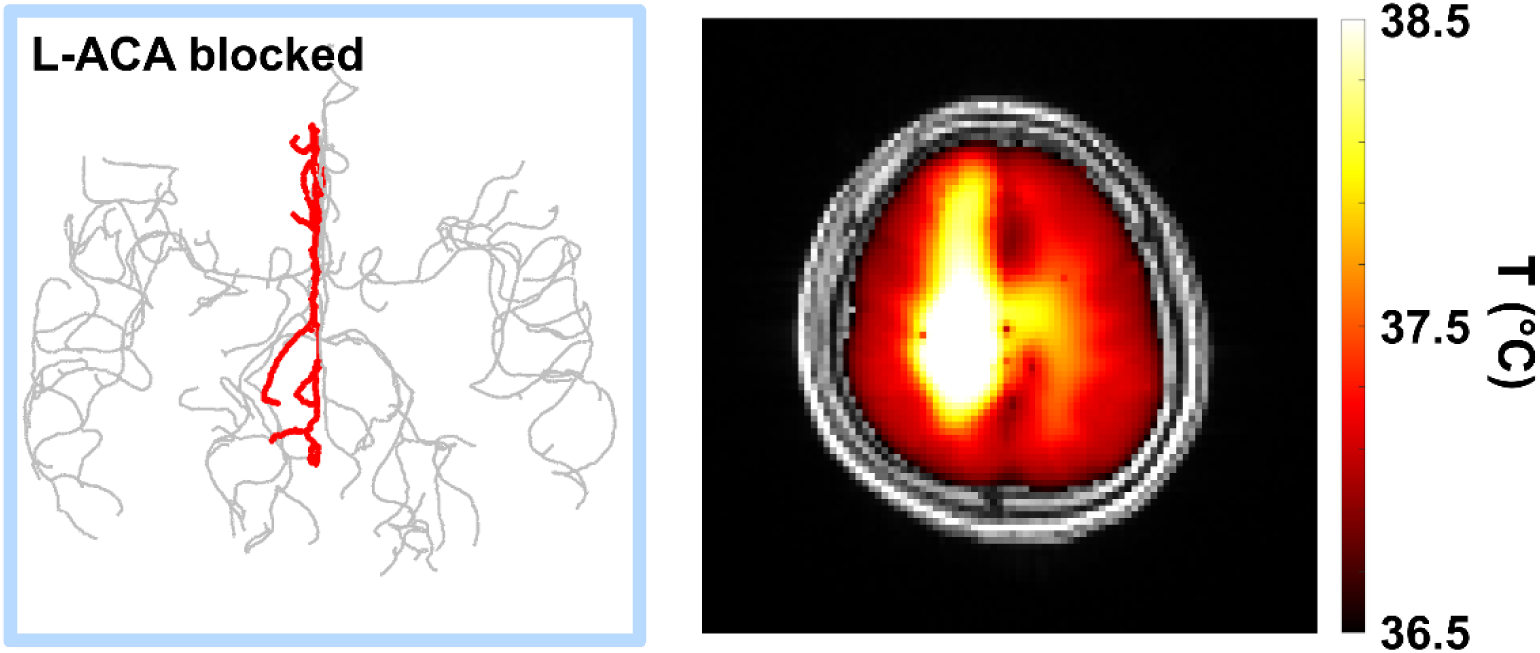
Temperature predictions after a simulated vessel occlusion using CVIG-derived vessel tree with left ACA blockage. The blood vessel occlusion scenario in which the left anterior cerebral artery (L-ACA) branch was selectively blocked in the CVIG-derived vessel tree. Red curves indicate the blocked vessel branches, and gray curves indicate the remaining vascular graph. Simulated brain temperature map under the L-ACA occlusion scenario. The localized thermal perturbation appears in the vascular territory corresponding to the blocked L-ACA branch, demonstrating that the topologically resolved CVIG graph enables side-specific and branch-specific temperature response.

**Equation S1.** To generate the arrival-time map for vessel tracking, we solved the Eikonal equation over the segmented vessel domain. In Eq. S1, *T_single_*_,*i*_(*x*, *y*, *z*) denotes the arrival-time from a given single inlet, and *F*(*x*, *y*, *z*) denotes the local propagation speed derived from the vessel-enhanced image or transformed vessel domain. The inlet was assigned a zero-arrival-time boundary condition, *T_single_*_,*i*_(*i^t^*^ℎ^ *inlet*) = 0. This formulation produces a scalar arrival-time-map in which vessel paths can be traced from distal high arrival-time regions toward proximal inlet-connected regions by following the descending direction of *T_single_*_,*i*_.

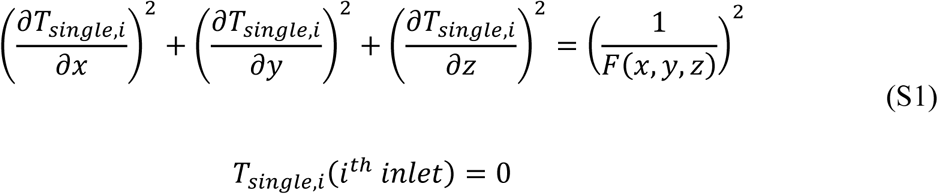

**Equation S2.** The multi-inlet arrival-time map was generated by taking the voxel-wise minimum across all single-inlet arrival-time maps. Here, *T_multi_*(*x*, *y*, *z*) denotes the final multi-inlet arrival time used for vessel tracking, and *T_single_*_,*i*_(*x*, *y*, *z*) denotes the arrival-time map generated from the *i*^th^ inlet. For MRA processing, *i* = 3, for MRV processing, *i* = 2. This operation assigns each voxel to the closest inlet in terms of cumulative propagation cost, thereby producing a unified guidance map for multi-inlet vessel tracking.

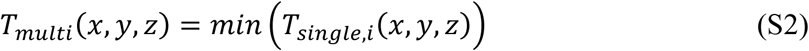

**Equation S3.** Centerline coverage was used to quantify the degree of completeness in recovering the reference vessel structure by a reconstructed vascular graph. In Eq. S2, *S_L_* denotes the reference centerline set or total reference centerline length, and *S_P_* denotes the predicted centerline set from a reconstruction pipeline. The intersection *S_L_* ∩ *S_P_* represents the portion of the reference centerline recovered by the prediction. Thus, coverage measures the fraction of reference vessel length captured by the reconstructed graph, with higher values indicating improved vascular coverage.

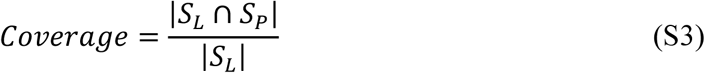

